# Intracellular Cytomatrix, Immobilized Biocatalysis, Matrix Micromechanics and The Warburg Effect: Entanglement of Two Age-Old Mysteries of the Normal and Malignant Cell

**DOI:** 10.1101/2023.04.07.535779

**Authors:** Tattym E. Shaiken, Mohamad Siam, Joel M. Sederstrom, Padmini Narayanan

## Abstract

For decades cancer studies have focused on molecular genetics while the role of the cytoplasm has remained obscure. Separation of the viscous fluid cytosol and elastic-solid cytomatrix has offered an opportunity to solve an age-old mystery in biochemistry; how millions of complex chemical reactions can occur simultaneously within the cell cytoplasm. The cytomatrix contains structural proteins, ribosomes, and metabolome enzymes responsible for unique biosynthetic pathways that involve immobilized biocatalysis. Immobilizing these catalytic complexes overcomes the spatial limitations for biochemical processes and allows integration of the intracellular and extracellular matrices and receptors with nuclear processes. Together, the cytosol and cytomatrix produce an interconnected synergistic network that maintains the operational flexibility of healthy cells as well as the survival of malignant cells. The cytomatrix is also responsible for cellular micromechanics and cytoplasmic motion. The combination of mechanical and biocatalytic processes triggered by extracellular signals and gene mutations in malignant cells requires additional energy. Cancer cells, consequently, utilize aerobic glycolysis, the Warburg effect, to meet the energy demands of the matrix mechanics that arise in response to imbalanced signaling and excessive biocatalytic activity. Clinical cancer is a rare event despite a high frequency of mutations, as clinical cancer is limited by the requirement for alterations that result in a high energy production state. Without these transformations, potential cancers can only survive in the quiescent state or will be eliminated. Survival of cancer cells indicates that the cancer cells were able to synchronize energy output for matrix mechanics supplying sufficient energy for tumor growth. Thus, the Warburg effect connects genetic aberrations and intracellular matrix mechanics with the ability to provide the energy supply required for the unprecedented complexity of tumor growth.

## Introduction

The somatic mutation theory of cancer considers malignant tumors as a consequence of sequential mutations of multiple genes. Genome-wide sequencing has identified many altered genes and epigenetic changes within single cancer cells (Tomasetti et al., 2015). It is unclear how many and which types of mutations are required for a normal cell to become a cancer cell. In cancer cells, alterations of the epigenome often disable DNA repair functions making cancer genetics and epigenetics two sides of the same coin (You and Jones, 2012). It is currently thought that constituent genetic and epigenetic alterations in malignant tumors are responsible for cancer cell survival such that the “fittest” cells survive because genetic and epigenetic alterations are able to rewire critical metabolic pathways. Cancer genes are common, as genetic studies of more than 28,000 tumors from 66 cancers revealed 568 cancer genes (Martinez-Jimenez et al., 2020). Despite the extensive detailed study of cancer genes and gene mutations, this approach has yet to provide a clear understanding of malignant cell survival.

A role for the cytoplasm is largely missing from current cancer theory. Genes without the presence of a cell’s cytoplasm are lifeless and have been described as a lonely library with no one to read the books within it (Mukherjee, 2022). The cytoplasm of living cells is a dynamic environment, and the motion arising from fluctuating forces was considered the result of thermally-induced random diffusion. Recently, using force spectrum microscopy (FSM), it has shown that the ubiquitous fluctuating motion in cells is not thermally induced (Guo et al., 2014). From the physicochemical standpoint, the cytoplasm is a two-phase system that can be separated into an elastic solid (cytomatrix) and viscous fluid (cytosol). We previously provided evidence of compartmentalization of transcriptome, proteome, metabolome, and protein synthesis within the cytomatrix and cytosol (Shaiken et al., 2023). The cytomatrix and extracellular matrix are interconnected, forming a singular integrated system with the receptors and signaling pathways. This two-phase system potentially can explain one of the oldest problems of cell biology; how diverse biochemical reactions occur within a viscous environment.

Here we examined the proteome profile of the solid phase, cytomatrix, using mas-spectrometry and extracellular and intracellular signaling pathways using the Hallmark and KEGG database to elucidate a key property of cancer cells, the Warburg effect. We propose that the genetic mutations that trigger protein overexpression (for example, HER2) and epigenetic alterations cause imbalanced signaling that requires an additional energy source for cytomatrix motion. As in the muscles of a fast-running athlete, the glycolytic pathway converts glucose into lactate to provide energy, in addition to mitochondrial energy production in cancer cells, to increase intracellular motion, intensify biocatalysis, and signal transduction within the viscous environment of the cytoplasm.

## Results

### Mass-spectrometry profile of the cytomatrix proteome

Applying different salt solutions, the nuclear-associated fraction was separated from the cytosol and nucleus (Shaiken and Opekun, 2014). Further, high-throughput methods such as RNA-seq, Ribo-seq, and mass–spectrometry were used to define the nuclear-associated fraction as an elastic solid phase of the cytoplasm – the cytomatrix (Shaiken et al., 2023).

The peptide area-based quantification method, the iBAQ (intensity Based Absolute Quantification), was used to identify the proteome spectrum of the cytomatrix. The protein abundance was calculated on the median normalized iBAQ that revealed the cytomatrix proteome profile. The structure-forming elements were investigated to determine the apparent contribution of the cytoskeletal and organelle-forming proteome in the solid phase assembly (Table S1). The proteome was categorized according to functional identity and complex assembly and the structure-forming proteome was divided by high and low numbers of the group members (Fig. 1A and 1B). Motor proteins such as actin, myosin, and tubulin and their binding proteins were the most abundant groups of the cytomatrix proteome. Actions created by these proteome groups, along with the dynein/kinesin and mobile keratin (Windoffer et al., 2011) are likely responsible for intracellular movement (Guo et al., 2014). Apparently, the intercellular contacts may orchestrate the actomyosin dynamics (Arnold et al., 2017). The next most abundant groups of proteins were transmembrane proteins and the ER-Golgi proteome, which suggests that membrane-forming structures such as ER, Golgi, and plasma membrane are integral parts of the cytomatrix. This hypothesis is supported by GOBP, KEGG, and Reactome pathway analyses (Shaiken et al., 2023). A nuclear connection to the cytomatrix is indicated by the presence of SUN1/2, emerin, and plectin, along with the nucleoporins. The presence of extracellular tight junctions, desmosomes, integrins, ezrins, and adhesion molecules show that the intracellular matrix is connected to the extracellular matrix and is able to play an essential role in intercellular signaling.

**Figure 1.**
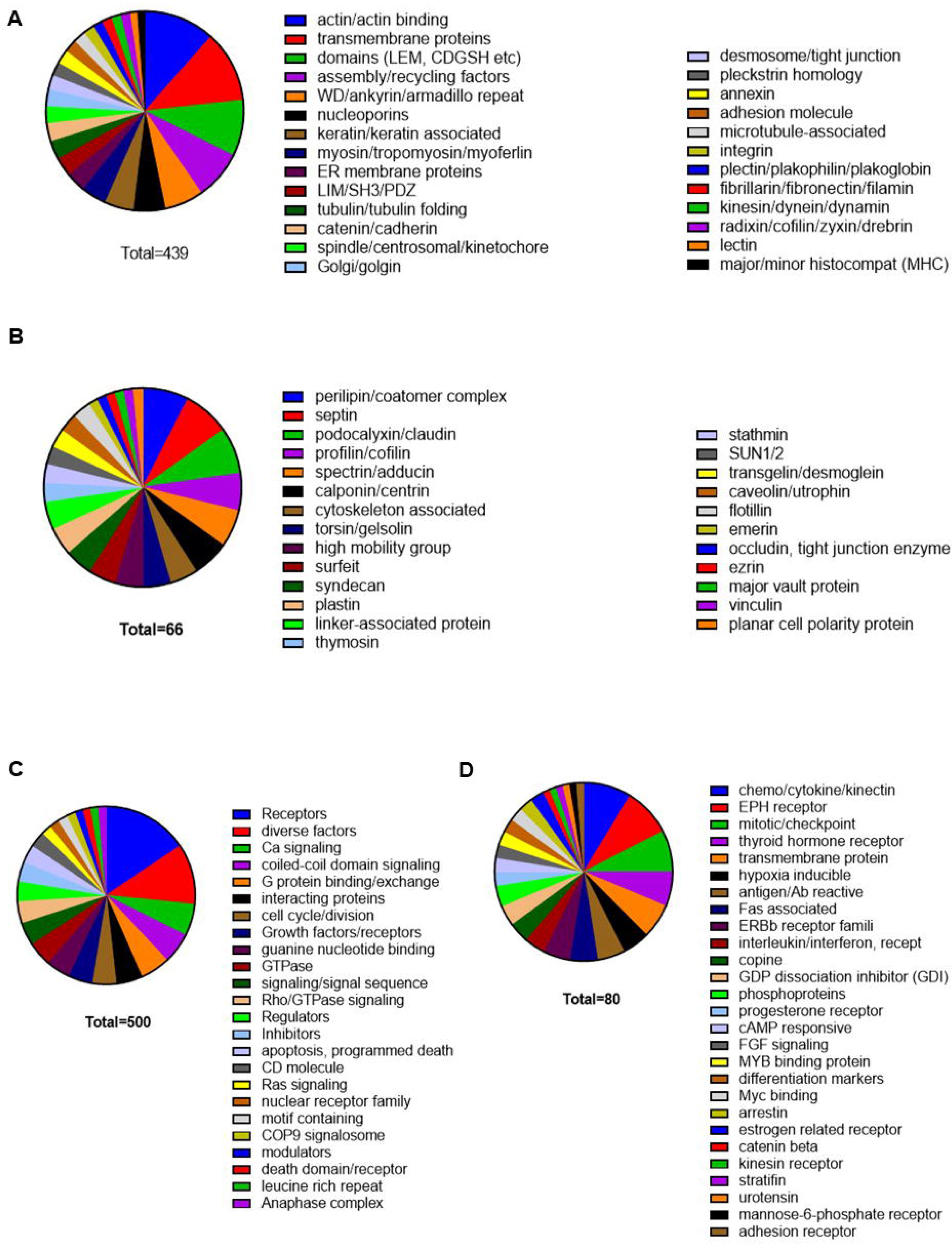
Pie charts of the HCT-15 cytomatrix proteome. Structure forming proteome (A) with the high member numbers and (B) with the low member numbers within the group. Receptors and signal transduction elements (C) with the high member numbers and (D) with the low member numbers within the group.

The cytomatrix apparently plays an essential role in signal transduction through extracellular and intracellular receptors and signaling molecules (Fig. 1C and 1D, Table S2). Diverse growth factor receptors, along with the <u>T</u>yrosine <u>K</u>inase <u>R</u>eceptors (TRK) and their downstream effectors, are also abundant within the cytomatrix. Cell division cycle and apoptotic pathways, diverse types of signal regulators, modulators and inhibitors, signaling sequences and signalosomes, Ca, cAMP, and G-protein signals were also abundant; they can deliver the normal or malignant functional requirements of the cell.

Oncogenic signals can be transduced through proto-oncoproteins and TRK (Fig. 2A and Table S3). The spectrum of enzymes shown in Figure 2B and Table S4 supports a multiplicity of chemical reactions, which suggests that the metabolic responses that could exclude each other coincide. Moreover, a host of kinases and phosphatases (Fig. 2C and 2D, Table S5) phosphorylate and dephosphorylate various specific substrates and modulate the activities of the proteins together in response to external stimuli. The question remains how these diverse chemical reactions can occur simultaneously in such a tiny gel-like environment, which has been broadly acknowledged for over a century (Taylor, 1923).

**Figure 2.**
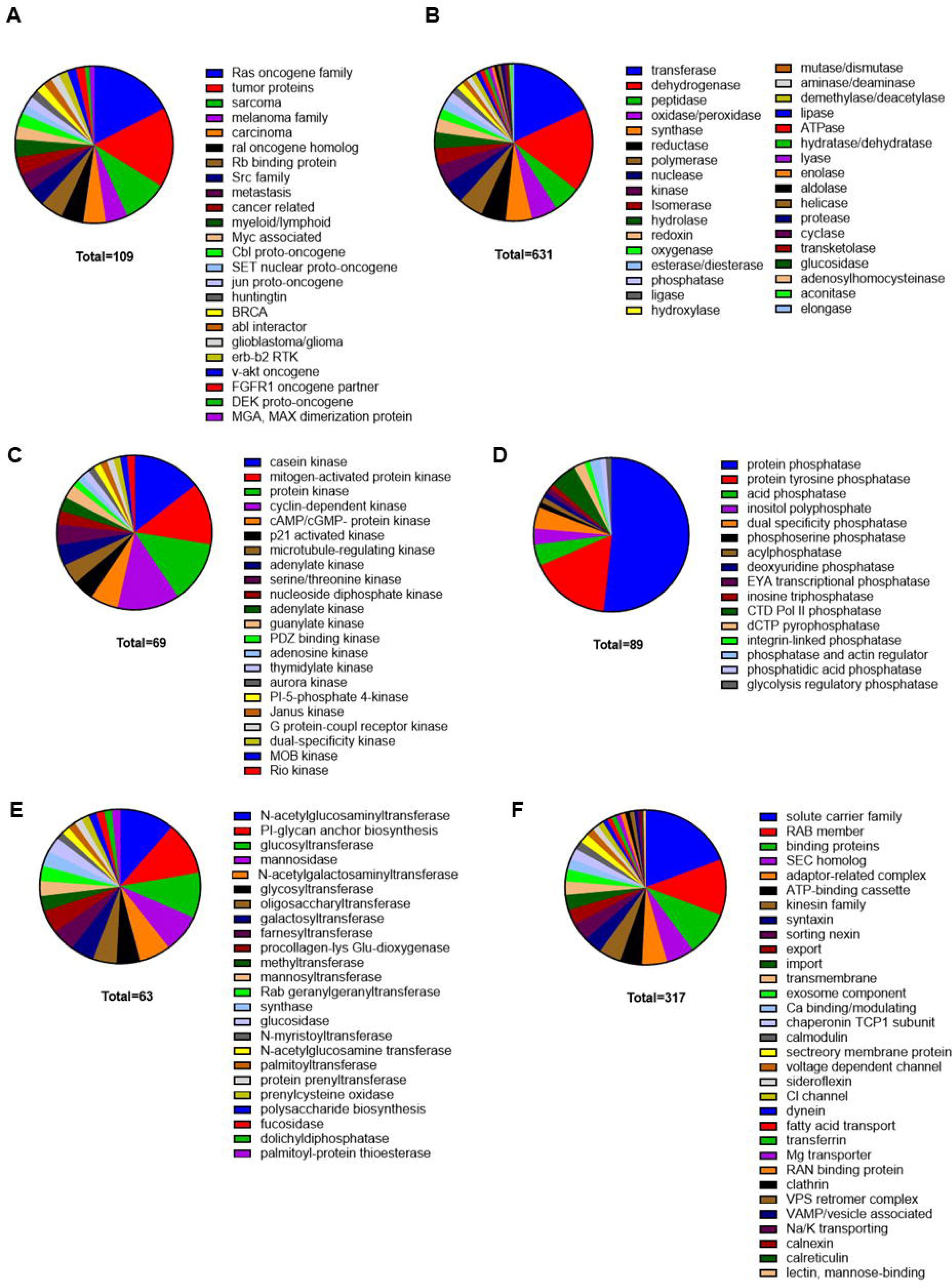
Pie charts of the HCT-15 cytomatrix proteome. (A) Proto-oncogenic signal transduction elements. (B) Enzyme spectrum. (C) Kinases. (D) Phosphatases. (E) Enzymes of the protein post-translational modification. (F) Transporters and intracellular traffic proteins.

Current knowledge shows several physicochemical problems of a gel-like cytoplasm. Although the cytoplasm is crowded, the individual molecule concentration is low, which conflicts with the principles of the mass action law. Furthermore, coinciding synthesis and degradation reactions could exclude each other within a limited space. Spatial impediments of the gel-like structure could also block free diffusion and the viscosity of the cytoplasm diminishes Brownian movement, which may slow reaction rates. Because of the limitations of a gel-like environment, the mechanisms of chemical reaction in the cell are routinely explained based on the idea of free diffusion as if the cells contained an aqueous solution, which would render the gel concept groundless (Pollack, 2001).

Co-isolation of the solid elastic component of the cell cytoplasm along with the enzymes that both catalyze diverse metabolic processes and regulate the activity of proteins, including the cytomatrix proteins, implies the presence of immobilized biocatalysis on the cytomatrix platform. Immobilization of metabolic pathways allows for the separation of chemical reactions and efficient biocatalysis. Immobilization and segregation can thus surmount spatial impediments to chemical reactions. Activities of the molecular motors (Guo et al., 2014) and actomyosin dynamics (Arnold et al., 2017) can also produce highly directed motions and cytoplasmic fluctuations, forming a ripple effect that overcomes the limitations of the gel-like structure and molecular crowding. The concept that cellular operation involves the directed mobility of the cytosolic substances through the intracellular matrix activity induced by the extracellular stimuli better describes cellular micromechanics. It explains how diverse metabolic reactions could coincide with immobilized catalytic complexes.

The most abundant and diverse post-translational protein modification is the attachment of carbohydrate and lipid moieties to proteins, and alterations have been shown to correlate with the progression of cancer and other disease states (Kobata and Amano, 2005; Zhao et al., 2009). The analysis of lipid and carbohydrate synthesis and enzymes of sugar/lipid post-translational modification of proteins revealed that the cytomatrix is the essential compartment for these processes (Table S6). Examples of enzymes that synthesize and attach diverse types of oligosaccharides, polysaccharides, glycerolipids, and isoprenoids/lipid moieties to proteins and glycoproteins through the nitrogen atoms of asparagine, O-linked glycosylation or farnesylation/geranyl-geranylation and palmitoylation through the thiol groups of cysteine or serine/threonine residues of proteins, are shown in Figure 2E. These sugar and lipid metabolism enzymes are critical in the biosynthetic-secretory function of the ER and Golgi apparatus (Reily et al., 2019). Moreover, the presence of these enzymes implies that the ER and Golgi are an integral part of the cytomatrix, which is also supported by the structural proteome analysis (Fig. 1 and Table S1). Consequently, the cytomatrix organizes Golgi-ER pathways into a singular integrated transport system (Fig. 2F and Table S7). The SLC and ABC transporters, along with the Rab vesicle transporters, are the most abundant groups within the cytomatrix. The list of transport systems supports the notion that elastic solid intracellular cytomatrix consists of the cytoskeleton, well-organized enzymatic clusters, and signaling complexes, which provide intracellular and extracellular traffic of molecules.

To distinguish the cytomatrix proteome from the cytosolic proteome and vice versa, we calculated the abundance ratio by dividing scores of iBAQ Normalized Medians of corresponding compartments to determine the fold differences. About 1,000 proteins of the cytomatrix and 500 of the cytosolic proteins are shown in Tables S8 and S9. We selected those proteomes that have at least 10-fold differences. For example, SUN1 (Sad1 and UNC84 domain containing 1) showed 10282-and 630-times higher values in the cytomatrix than in the cytosol in the duplicate analysis. Thus, the proteome profile of the cytomatrix and cytosol indicates the presence of two distinct phases, the elastic solid intracellular matrix, and viscous fluid cytosol within the cytoplasm, that should be considered complementary to one other and play an essential role in the fine-tuning of biochemical processes. Thus, for the proper functioning of the cell containing an elastic solid cytomatrix with immobilized catalytic complexes and the cytosol–viscous fluid, the cytoplasm must be in constant motion (i.e., a condition that requires energy expenditure). The mechanical part of the motion can rely on the cytomatrix because it forms the complex solid system of intracellular and extracellular matrices and receptors. Receptor-ligand binding can trigger cytomatrix motion that plays the role of the pump for the cytosol. Cancer cells which characteristically have an overproduced proteome, elevated ribosome biogenesis (nucleolar hypertrophy), signaling systems, and metabolome imbalance, require more energy than can be provided by mitochondria. This extra energy could be delivered by glycolysis and is expressed as the Warburg effect.

### Ribosome footprint analysis of the cytomatrix

Ribo-seq was used to identify differentially expressed genes that permit over-representation (or enrichment) analysis (ORA), a statistical method that determines whether genes from pre-defined sets are present more than expected. Here, the KEGG collection of databases and Hallmark gene sets were used to examine the potential role of cytomatrix pathways in cancer development. The KEGG pathway is a collection of manually drawn pathway maps representing networks for signal transduction, metabolism, cellular processes, and human diseases, including cancer (Chen et al., 2020). The over-representation analysis of the cytomatrix over cytosol was performed for the cytomatrix dataset and normal growth condition over the mTOR kinase inhibitor (TOR-KI) pp242 – for the cytomatrix and cytosol datasets, respectively. Of note, TOR-KI inhibits mTOR-dependent protein translation and directly affects autophagy (Abu-Remaileh et al., 2017; Liu and Sabatini, 2020).

We compared the upregulated (Fig. 3A and 3B) and downregulated (Fig. 3C and 3D) cytomatrix pathways of a colon cancer cell line cultured at the normal growth conditions (cytomatrix over cytosol) with the cytomatrix (cytomatrix over cytomatrix pp242) and cytosolic (cytosol over cytosol pp242) KEGG pathways of the HCT-15 cells treated with pp242 (Table S10). The exclusive localization of the calcium and phosphatidylinositol signaling pathways (Fig. 3A) confirms the notion that the ER and PM are integral parts of the cytomatrix. The pathways in cancer, TGF beta, WNT, Hedgehog, and Notch signaling pathways were also upregulated in the cytomatrix but downregulated in the cytosol and within the cytomatrix treated with pp242. Several other pathways were downregulated within the cytomatrix, such as immune response and tumor formation signaling pathways: Jack/STAT, Nod, Rig I, and Toll-like receptor signaling. Non-receptor signaling pathways such as Adherens, Focal adhesion, Gap, and Tight junction that can orchestrate the signaling output through actomyosin dynamics (Arnold et al., 2017) were upregulated in the cytomatrix of cells cultured under normal growth conditions but downregulated in the cytosol and in the cytomatrix after the treatment with the pp242 (Fig. 3B). These results show that the upregulated pathways of the cytomatrix are in an inverse relationship with the downregulated same pathways of the cytosol and the cytomatrix with the pp242-treated cells.

**Figure 3.**
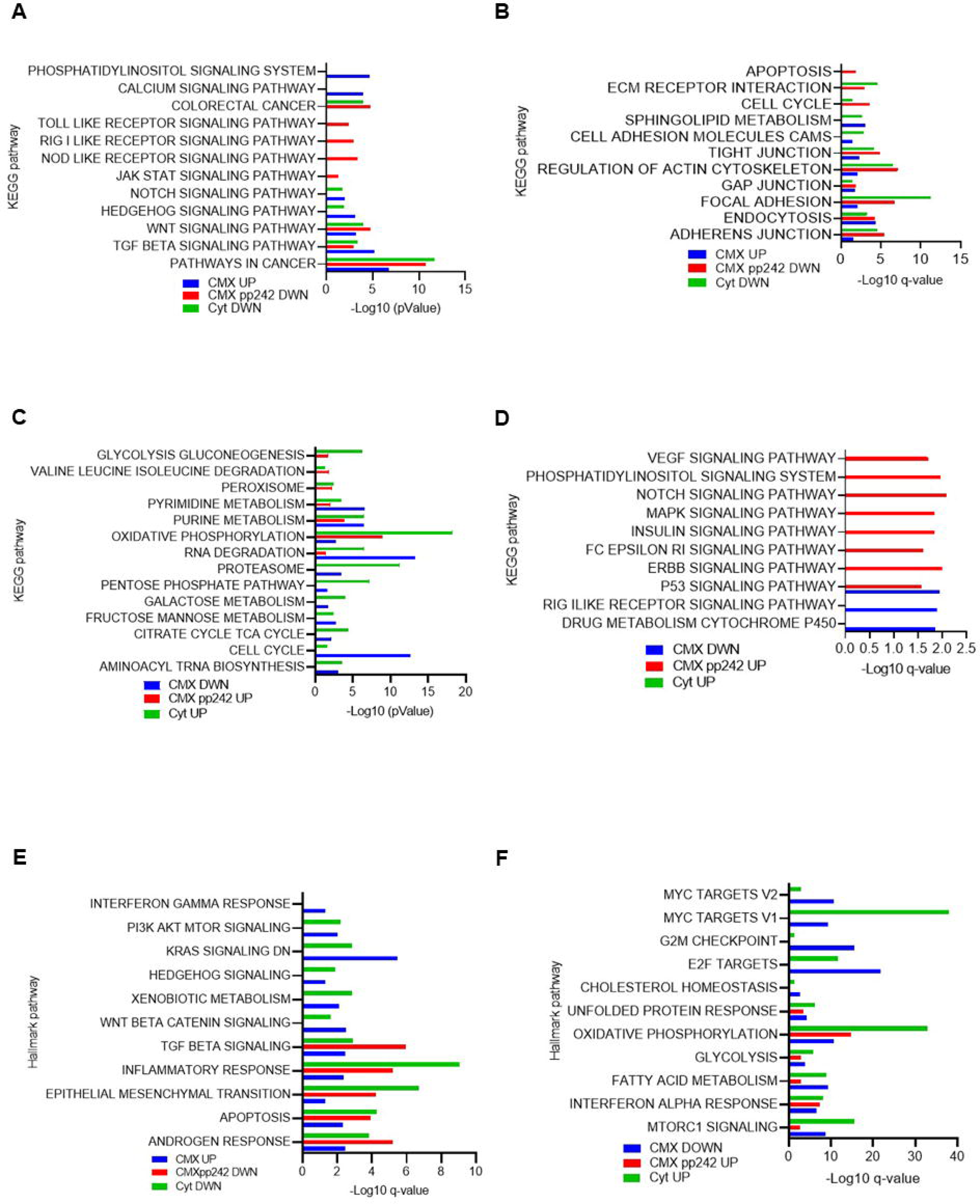
Upregulated and downregulated selected pathways of the cytosol and cytomatrix of HCT-15 cells identified through Ribo-seq. (A - D) Categorized KEGG pathways. (E, F) Categorized Hallmark pathways

Examining the downregulated pathways of the cytomatrix revealed a different spectrum of the gene sets that had an inverse relationship with the cytosol where those pathways were upregulated (Fig. 3C). These pathways include tRNA biosynthesis, cell cycle, TCA cycle, carbohydrate metabolism, and protein degradation by the proteosome. The RNA degradation, oxidative phosphorylation, purine and pyrimidine metabolism, amino acid degradation, and glycolysis/gluconeogenesis pathways were upregulated in the cytosol, and TOR-Ki treated cells cytomatrix but downregulated within the cytomatrix cultured under normal growth conditions. Thus, TOR-Ki treated cells show parallel upregulation of the energy generation and metabolic pathways in the cytosol and the cytomatrix, consistent with the role of the cytomatrix in cell survival under environmental alterations.

Several other pathways were upregulated, such as inflammatory signal (FcεRI), Insulin, Notch, VEGF, MAPK, ErBb, and phosphatidylinositol signaling pathways revealed by the cell cytomatrix treated with the TOR-KI (Fig. 3D). Upregulation of p53 pathways corresponded with the downregulation in the cytomatrix of cells cultured under normal growth conditions, where we also observed the downregulation of the drug metabolism and Retinoic acid-inducible gene set (RIG I like receptor). None of these pathways were detected in the cytosol.

Hallmark gene sets represent specific, well-defined biological states and provide more refined and concise inputs for gene set enrichment analysis (Liberzon et al., 2015). Hallmark gene sets may define the role of the extracellular and intracellular signaling in cytomatrix micromechanics. The cell cytomatrix cultured at the normal growth condition revealed a wide range of extracellular and intracellular signaling pathways (Fig. 3E and Table S11). The upregulated signaling pathways were categorized as inflammatory and survival pathways (IFNg, IL2, TNFa, apoptosis, and PI3K-Akt-mTOR signaling) and tumor-promoting pathways (Androgen, TGFb, KRAS, WNT, Hedgehog, and EMT signaling), which were downregulated in the cytosol and within the cytomatrix of cells treated with pp242.

The energy generation, protein synthesis, and degradation as well as ribosome biogenesis pathways were downregulated in the cytomatrix; however, they were upregulated regarding pp242 treated cell cytomatrix (Fig. 3F and Table S11).

ORA of the cytomatrix over cytosol reveals the enriched pathways of the cytomatrix and ORA cytomatrix over cytomatrix of the cells treated with the TOR-Ki exposed drug-treated cytomatrix pathways. Thus, upregulated pathways disclosed with the pp242 treatment and those not exposed at the cytomatrix over cytosol are the results of reprogrammed expression as shown in Figs. 3D and 3F, which include signaling pathways, energy, and metabolic pathways (red lines).

The overexpression of the signaling pathways in cancer cells is the cause of high metabolic activity in the cytoplasm. Cancer therapy, on the other hand, eliminating highly metabolically active cells, possibly promoting the survival of normalized ones, may endorse cancer recurrence by metabolic reprogramming, similar to TOR-KI treated cell lines.

### Robustness of the cytomatrix permits isolation of malignant cell from frozen tumors

At super-low temperatures, the biochemical processes of the cell are halted, and tumors can be stored indefinitely. Tumors stored at –80°C maintain tissue morphology, and long-term, low-temperature storage did not adversely affect the quality of RNA (Andreasson et al., 2013; Grizzle et al., 2010).

Revealing the cytomatrix allowed to develop of a cell extraction method using deep-frozen (–80°C or liquid nitrogen) tissue samples and permitted the separation of authentic malignant cells from the tumor stroma (Fig. 4A). Malignant cells were isolated from flash-frozen mouse osteosarcoma (OS) tumors and stained with the hematoxylin-eosin (H&E, Fig. 4B) and the silver (Ag)-based staining of nuclear organizer regions (AgNor stain, Fig. 4C). The AgNor staining of extracted cells showed multiple nucleoli (nucleolar hypertrophy) within each cell nucleus, which is a feature of malignancy. Patient-derived xenografted (PDX) Ewing sarcoma tumor cells stored at –80°C were also isolated and stained with the H&E (Fig. 4D). Because Ewing sarcoma expresses CD99 (Dubois et al., 2010), cells can be detected by flow cytometry. We used multicolor flow cytometry to distinguish Ewing sarcoma cells (A673 and TC71) from mouse cells of hematopoietic origin (CD45), and macrophages (CD14). Ewing sarcoma cells A673 and TC71 were isolated from Mouse xenograft tumors stored for several years at −80°C. Extracted cells were stained with corresponding fluorochrome conjugated antibodies (Table 1, Fig. S1). Results show that cell surface antigens were detectable and intact and cells sorted by cell surface antigens.

**Figure 4.**
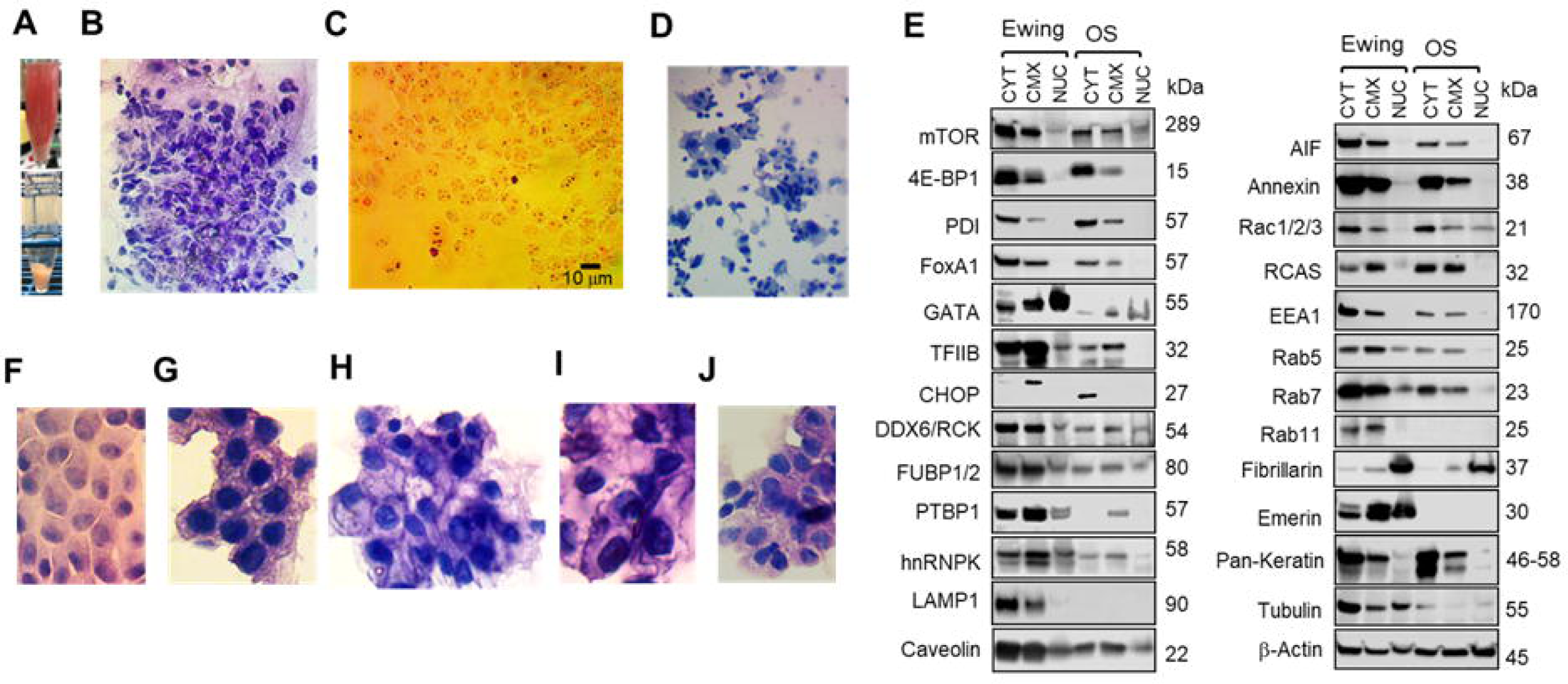
Morphological and biochemical analysis of malignant cells isolated from flash-frozen tumors. (A) Snapshot of malignant cell isolation from tumor stroma. Osteosarcoma cells isolated from the mouse tumors H&E staining, (C) AgNOR staining. (D) Malignant cells isolated from the patient-derived xenograft (PDX) Ewing sarcoma tumors, H&E staining. (E) Western blots of the osteosarcoma and Ewing sarcoma cells isolated from frozen tumors. Cells from human tumors (F) Lung, (G) Prostate, (H) Pancreatic, (I) Kidney, and (J) Stomach cancer cells. 100x, oil.

**Table 1.** Flow cytometry analysis of Ewing sarcoma.

| Isolated Ewing sarcoma cells from frozen tumors | A673 | TC71 |
| --- | --- | --- |
| CD99 positive cells from Total cell included in SSC single cells gate | 66.1 % | 32.1 % |
| CD45 positive cells from Total cell included in SSC single cells gate | 0.66 % | 0.59 % |
| CD45 positive/CD14 Positive cells from Total cell included in SSC single cells gate | 0.14 % | 0.22 % |
| CD45 positive/CD14 negative cells from Total cell included in SSC single cells gate | 0.52 % | 0.37 % |

We then compared protein expression in OS and Ewing sarcoma in isolated tumor cells using western blotting (Fig. 4E). Although some proteins were not detected due to antibody specificity for human (Ewing) and mouse (OS) proteins, the fractionation results indicated that cells could be biochemically characterized.

Malignant cells were also isolated from tumor specimens (Biooptions, Brea, CA) and stained with the H&E (Fig. 4 F-J). Results show that individual cancer cells can be isolated from tumors stored in deep-frozen states for decades. Isolation of cancer cells from frozen tumors and detection by surface antigens show that intracellular and extracellular matrices are robust and integrated structures.

## Discussion

Depending on the organ or tissue, the energy sources may vary with the needs of the cell, which ultimately determine the amount of energy consumed. In healthy eukaryotic cells, oxidative phosphorylation (oxphos) is the primary energy extraction pathway. The most energy consuming cells are muscle cells during exercise and growing malignant cells. Whereas fat oxidation is essential at lower intensities of energy needs, carbohydrate oxidation dominates at higher exercise intensities and is associated with progressive increases in glucose uptake coinciding with the workload (Watt et al., 2002). Skeletal muscle contractions stimulate glucose uptake during exercise and involve complex molecular signaling processes distinct from the insulin-activated pathway. Exercise increases the uptake of glucose up to 50-fold to ensure the maintenance of muscle energy supply during physical activity (Sylow et al., 2017). Almost every regulatory aspect of carbohydrate metabolism is designed for the rapid provision of ATP (Miller et al., 2002). As exercise intensity increases, the mitochondria are unable to oxidize all of the available pyruvate resulting in increasing pyruvate concentration that triggers the conversion of pyruvate to lactate via the enzyme LDHA, lactate dehydrogenase A (van Hall, 2010).

LDHA is aberrantly highly expressed in many cancers and is associated with malignant progression (Feng et al., 2018). It is estimated that tumor cells utilize up to 30 times more glucose than normal cells and produce up to 43 times more lactic acid, irrespective of oxygen supply (Holm et al., 1995). In cancer cells, high glucose uptake drives an insulin-independent proliferative phenotype (Bollig-Fischer et al., 2011), similar to muscle cells. Thus, in malignant cells, as in muscle cells, the glucose uptake and lactate production mechanisms are dictated by the energy demand. The Warburg effect in malignant cells involves enhanced lactate production that results from mitochondria being unable to oxidize all the available pyruvate.

The triggering of the Warburg effect is a complex process that involves multiple regulators. In tumorigenesis, overproduced or mutated growth factors activate transcription factors HIF-1a, NFkB, and c-Myc via the RTKs-PI3K-Akt-mTOR pathway, leading to the expression of glucose transporters and glycolytic enzymes (Fan et al., 2010; Yu et al., 2017). The overexpression of GLUT1, the insulin-independent transporter (Ebeling et al., 1998), also contributes to the increase of glucose uptake, lactate production, resulting in cancer cell survival. The overexpression of hexokinase 2 and other glycolysis enzymes also contributes to the enhanced glycolytic rate and promotes tumor progression (Nowak et al., 2018). The glycolytic pathway is adjusted in response to cell conditions. The factors that regulate glycolysis do so primarily to provide adequate amounts of ATP to meet cellular demands (Fig. 5 A-C). However, malignant growth disrupts glycolytic regulation (Fig. 5D). The insulin-independent glucose uptake, overexpression of rate-limiting glycolytic enzymes, and lactate excretion are potential ways to overreach the energy threshold that controls the glycolytic pathway in normal conditions.

**Figure 5.**
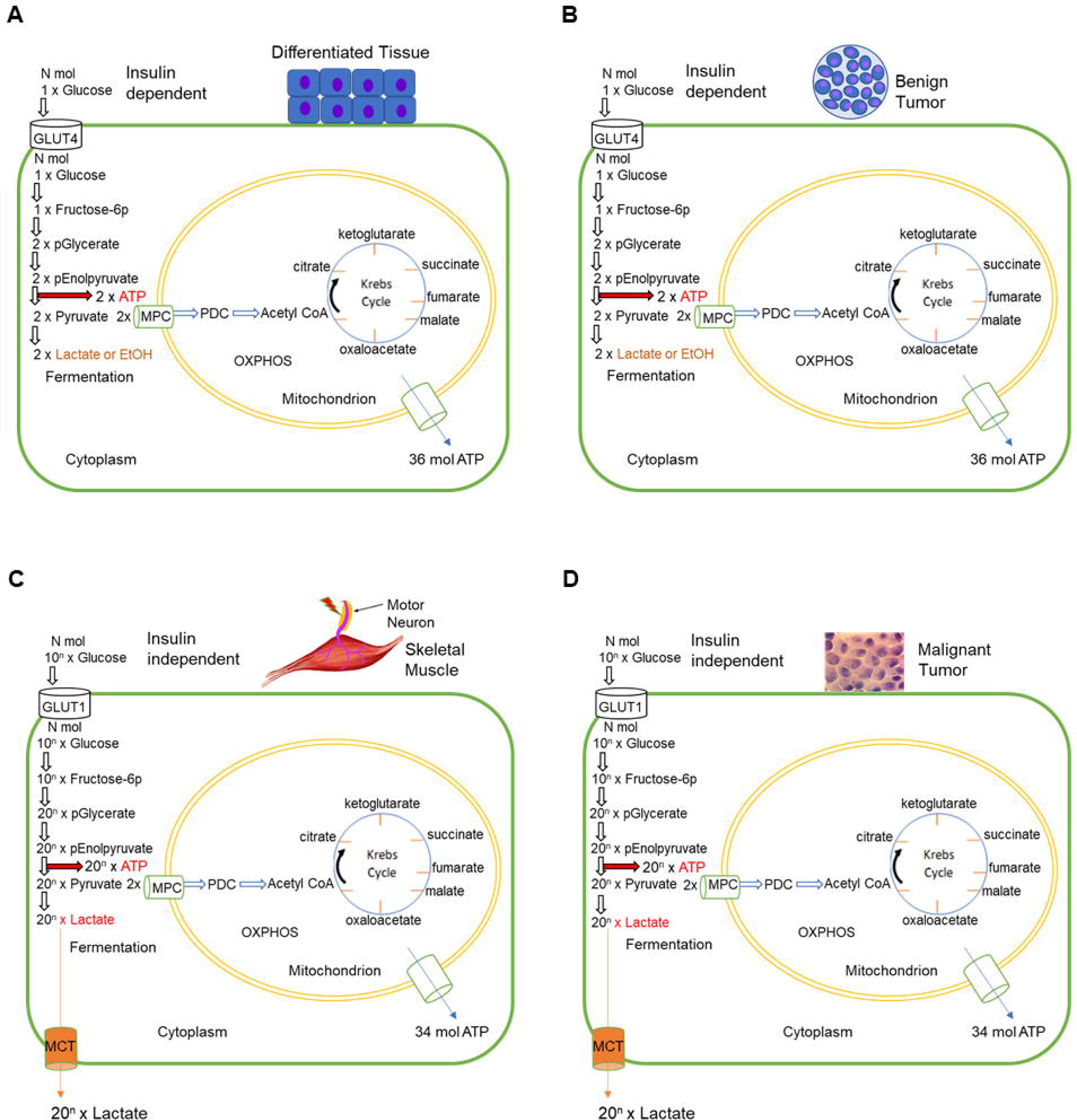
Differences and similarities of healthy and malignant cell energy generation and consumption. In (A) differentiated tissue and (B) benign tumors the glucose uptake and ATP generation pathways are similar. The energy consumption in (C) high-workload muscle and (D) tumors is similar. The glycolytic pathway regulation is different in high workload muscle and tumors. Insulin and glucose transporters regulate glucose uptake in healthy, muscle, and benign tumor cells; cellular energy production is disrupted in malignant tumors. MPC – mitochondrial pyruvate carrier; MCT – monocarboxylate transporter.

On the other hand, a role of oxphos in cancer maintenance can not be ruled out. Interestingly, glycolytically suppressed cells upregulate mitochondrial function to compensate energy requirements of cancer cells (Shiratori et al., 2019). Restoration of growth of mtDNA-depleted tumors is associated with the horizontal transfer of mitochondrial genomes from host tissue and repair of respiration, which point to the importance of mitochondrial function to tumor growth and energy supply (Tan et al., 2015; Wallace, 2012). Therefore, cancer cell metabolism is diverse, with varying reliance on glycolysis and mitochondrial oxphos, as has been shown with LDH inhibition and knockout experiments (Fantin et al., 2006; Oshima et al., 2020). It remains unclear why cancer cells require energy beyond oxidative phosphorylation, where and what cellular process utilizes the surplus energy, and what mechanical process within the cancer cell requires such vast amounts of ATP.

Immobilization of biosynthetic pathways via the cytomatrix provides separation of the chemical reaction. For efficient biocatalysis, however, motion of the liquid phase of the cytoplasm is required. The force that moves the cytosol must be applied for the over 1 million chemical reactions per second that occur in a human cell (Kurien, 2004). The matrix system segregating biocatalysis performs efficient reactions and signaling in healthy tissues at normal conditions, primarily through mitochondrial respiration. It appears that, cancer cells often require more energy for the cytomatrix mechanics than do normal cells possibly due to the proteomic and signaling imbalances caused by the mutations and epigenetic alterations. As a consequence, the cytomatrix dynamics of cancer cells require a large amount of energy to support the high metabolic rate. The requirement for alterations in matrix mechanics to meet the biosynthetic and energy demands of malignant cells likely reflect the presence of critical transformations needed for the cancer to develop. Therefore, one might consider that both alterations in matrix mechanics as well as oncogenic mutations are essential for cancer development.

The solid phase of the cytoplasm, the cytomatrix, contains the entire cell as one robust mechanosensor unit. The cellular matrix is so stout that cells were isolated from frozen tumors stored for decades (Fig. 4F-J) without their receptors being degraded by long term storage at low temperatures as shown by the flow cytometry experiments (Table 1). A spectrum of the cytomatrix filamentous proteins was identified that could participate in the matrix motion (Fig. 1A and 1B) including the intracellular and extracellular signaling receptors and molecules (Fig. 1C and 1D) and proto-oncological signaling pathways (Fig. 2A) that could trigger and intensify the cytomatrix dynamics. We also confirmed the presence of a complex of catalytic enzymes (Fig. 2B), kinases (Fig. 2C), and phosphatases (Fig. 2 D) including the glycosylation enzymes (Fig. 2E) that participate in matrix forming along with members of the transport system (Fig. 2F) that participate in cytosolic motion. The proteome profile shows that the elastic solid cytomatrix has specific components for the active force, which results in cytoplasmic motion. The cytomatrix motion likely involves numerous and complex matrix filaments and cytoskeletal machinery producing a system comprised of, and regulated by, hundreds of dynamic proteins.

The most prominent types of dynamic filaments are actin microfilaments and microtubules, that act as tracks for movement of the molecular motors that convert chemical energy into mechanical energy. Kinesin and dynein motors both utilize microtubules for transport, whereas myosins are the only known actin-based motor proteins (Guo et al., 2014; Heissler and Sellers, 2016). We found that actin and actin-binding proteins were the most abundant along with the transmembrane proteins. We believe the actin-myosin fibers are likely the primary force in cellular and intracellular motility. Moreover, actin-myosin muscle fiber contraction triggers carbohydrate metabolism by the glycolytic pathway that leads glycogen breakdown to lactate during intense exercise (Hargreaves and Spriet, 2020), which is similar to the aerobic glycolysis (i.e. Warburg effect) seen in cancer cells.

Although there are specific differences between muscle and nonmuscle fiber content and regulation, the actin-myosin action could also largely be responsible for aerobic glycolysis. Actin represents 1–5% of all cell proteins in nonmuscle cells compared to up to 10% in muscle cells. The β-actin and γ-actin are the actin isoforms found mainly in non-muscle cells (Herman, 1993). Six actins, more than forty tropomyosins, fifteen classes of myosins and numerous other actin-binding proteins diversify and dynamically compartmentalize the mammalian actin cytoskeleton. Actin-binding proteins determine the speed of actin polymerization and the length of actin filaments. The monomeric globular (G)-actin, a 43-kDa ATPase, can self-assemble into filamentous (F)-actin. ATP hydrolysis in the filament is tightly coupled to polymerization and regulates the kinetics of assembly and disassembly, as well as the association of interacting proteins (Pollard and Borisy, 2003). The Cofilin, Formin homology (FH) proteins, and actin-related proteins complex, Arp2/3 complex, are major actin binding proteins that control nucleation, assembly and disassembly of actin filaments (F-actin). The Arp2/3 complex is regulated by its association with the WAVE and WASP family of WH2 domain containing proteins (Gautreau et al., 2022). The polymerization and depolymerization of F-actin regulate cell motility, cytoskeletal reorganization, also can promote epithelial-mesenchymal transition and metastasis (Goley and Welch, 2006; Olson and Sahai, 2009).

Upstream signals tightly regulate the activity of the actin polymerization machinery. Many of the regulatory factors are either membrane-anchored small G proteins of the Rho family or phospholipids (Olson and Sahai, 2009). Upon association with GTPases including RhoA, RhoB, Cdc42 or Rif the actin-binding FH proteins change conformation to promote actin polymerization. These complex regulatory mechanisms serve to ensure that actin is not polymerized inappropriately, and that actin polymerization can be increased in response to appropriate stimuli. Extracellular signals, including growth factors and extracellular matrix components, control the GTP-loading of Rac1 and Cdc42, the generation and hydrolysis of phospholipids, and the recruitment of adaptor protein complexes to membranes. In addition, many regulators of actin polymerization are phosphorylated in response to external stimuli; in some cases, this can have been shown to have very profound effects on their function. For instance, the Src kinase mediates growth factor signaling that causes oncogenic transformation, including dramatic changes in the actin cytoskeleton, cell shape, and motility (Tehrani et al., 2007). Moreover, the sheer number of actin post-translational modifications poses a serious challenge to understanding their regulatory mechanisms; how can an actin molecule, whose primary role is to generate dynamic filaments, when decorated with potentially structure-disturbing PTMs (Varland et al., 2019)?

The myosin superfamily is a large and diverse protein family involved in a number of cellular pathways. Myosins localize to a number of intracellular compartments, participate in many trafficking and anchoring events and organize the endomembrane system. The recruitment of motor proteins to an organelle or molecular complex is generally the result of the tail binding to a specific adaptor protein (Hartman and Spudich, 2012). A single mammalian non-muscle cell is well known to express multiple myosin isoforms that are specialized for particular cellular roles, with little in the way of overlapping functions (Peckham, 2016).

Actin-myosin complex play central roles in many biological functions such as cell motility, contraction, adhesion, cell division, junction formation, protrusion of the cellular membrane, vesicle trafficking, chromatin remodeling, and maintenance of the physical integrity of the cell (Biber et al., 2020). F-actin assembles with myosin II filaments to form a protein complex that uses energy from ATP hydrolysis to power actomyosin contraction (O’Connell et al., 2007). The motility of malignant cells has been intensely studied in cell migration experiments (Olson and Sahai, 2009). However, the actin dynamics in living tumors are different as the majority of cancer cells are not motile in vivo. Very few cells within primary tumors are motile; even in aggressive metastatic xenograft models, relatively few cells where shown to be motile (Sahai, 2007), which contradicts the results of the in vitro studies.

The question arising from the numerous regulatory steps of actin fiber assembly and actomyosin contractility is that if it has little effect on cell movements even within tumor cell motility, where is this activity applied, and why does it utilize aerobic glycolysis? We propose that the answer relates to the utilization of the active force for cytomatrix mechanics for the intracellular processes that require cytosolic movement to provide efficient biocatalysis. In this case, the extracellular receptors transmit signals to the regulatory factors, such as membrane-anchored small G proteins or phospholipids that activate actin-binding proteins to trigger the cytomatrix motion. This general scheme is outlined using KEGG and Hallmark pathway analysis. The KEGG ORA revealed that the extracellular signaling pathways consist of specific receptor groups (Fig. 3A) and intercellular connectors and channels through which cells communicate with each other (Fig. 3B). KEGG ORA also showed the flexibility of the extracellular and intracellular pathways during drug treatment (Fig. 3C and 3D). The Hallmark database provided another list of extracellular signaling (Fig. 3E) and flexibility of metabolic pathways during drug treatment, with the alternative signaling that emerges to reprogram metabolome (Fig. 3F). Thus, the mass-spec proteome and over-representative analyses of signaling pathways showed that the mechanical motions of the cytoplasm and biocatalytic processes are entangled to provide the efficient chemistry of the cell that exploits the energy sources from the mitochondria and glycolysis under both normal physiological conditions and malignant transformation. The cytoplasmic fluctuations are three times larger in malignant cells than in their benign counterparts (Guo et al., 2014); this could be explained by excessive mechanics imposed by imbalanced signaling and metabolic pathways that require an additional energy source and compensates by glycolysis, known as the Warburg effect.

Like muscle myofibrils of fast-running athletes, the malignant cell actomyosin that form contractile structures, along with the actin polymerization/depolymerization within the cytomatrix, consume the ATP from glycolysis that produces excessive lactate as a byproduct that efficiently extracts to the bloodstream avoiding feedback inhibition (Fig. 5C, D). Carbohydrate is the primary fuel for anaerobic and aerobic metabolism, whereas sprints (anaerobic ATP), as dictated by the game, are added to the contribution of aerobic ATP. As with Olympic caliber athletes, this scenario is repeated many times, with carbohydrates providing most of the aerobic and much of the anaerobic fuel (Hargreaves and Spriet, 2020) for cancer cells.

Cancer cells overexpressing growth factor receptors (for example HER2) intensify intracellular motion, which can increase the rate of chemical reactions in growing malignant cells. While anticancer drug treatment can eliminate the actively growing cells; cancer cells with slow intracellular motion requiring low glucose uptake consequently can maintain a rate of catalytic activity comparable to normal cells and thus survive anticancer therapy. Suppression of ChREBP (carbohydrate responsive element binding protein) in cancer cells results in diminished aerobic glycolysis, reduced glucose uptake and lactate production and stimulates mitochondrial respiration that results cell cycle arrest (Tong et al., 2009) suggesting that the switch from anaerobic to aerobic (or vice versa) ATP production for cellular mechanics in malignant cells may be one of the main reasons for cancer cell transformation and survival or elimination or the quiescent state. Clinical cancer is a rare event despite a high frequency of mutations, defined as passenger mutations, in the human genome sequence project (Greenman et al., 2007), as cancer is limited to only those mutations that result in the ability to produce a high energy production state. Thus, unless mutations lead to increased energy production, the potential cancer may only survive in the quiescent state (or benign) or it will be eliminated. Survival of cancer cells means that the cancer cells have acquired the ability to synchronize energy output to matrix mechanics so that there is sufficient energy for tumor growth (Video presentation).

As a concluding remark, the energy waste through Warburg effect can be one of the causes of cancer cachexia. Total energy consumption for the cytomatrix mechanics of cancer cells, which may include the actomyosin assembly and contractility, the dynamics of tubulin, keratin, and other fibrillar proteins, can be comparable to the dynamics of the athlete cell myofibrils during intense exercise according to glucose uptake and lactate production data. The Warburg effect connecting genetic aberrations, intracellular matrix mechanics, respiration, and glycolysis allows a better understanding of cancer physiology, which could be essential in fighting the disease.

Here, we demonstrated elements of a cellular engine, the cytomatrix, and suggested its role in the Warburg effect; the construction and operation of the whole machine remain to be discovered. We believe experimental approaches such as direct measurements of matrix mechanics can test our concepts in the very near future.

## Materials and Methods

### Cell lines and cell culture

Colorectal carcinoma HCT-15 (CCL-225, RPM-1640), cells were grown in the incubator under humidified conditions at 37°C under 5% CO2 in Dulbecco’s Modified Eagle Medium or RPMI-1640 supplemented with 10% (v/v) fetal bovine serum (FBS) and 1% (v/v) penicillin and streptomycin.

### Antibodies

Reagents were obtained from the following sources: antibodies to mTOR (2983), 4E-BP1 (9644), PDI (3501), FoxA1 (53528), GATA-6 (5851), TFIIB (4169), CHOP (2895), DDX6/RCK (9407), FUBP 1/2 (13398), PTBP1/2 (57246), hnRNPK (9081), LAMP1 (9091), Caveolin-1 (3267), AIF (5318), Annexin (32934), Rac1/2/3 (2465), RCAS1 (12290), EEA1 (3288), Rab5 (3547), Rab7 (9367), Rab11 (5589), Fibrillarin (2639), Emerin (5430), pan-Keratin (4545), β-Tubulin (2128), β-actin (3700) were purchased from Cell Signaling Technology (Danvers, MA); mTOR kinase inhibitor, TOR-KI, pp242 was purchased from Tocris Bioscience (4257/5).

### Antibodies for Flow Cytometry

| Laser | EM Filter | Marker | Color / Format | Host / Target | Company | Catalog Number |
| --- | --- | --- | --- | --- | --- | --- |
| 355 | 652LP | CD99 | BUV661 | Mouse anti-Human | BD Biosciences | 750427 |
| 355 | 450/50 | CD45 | BUV395 | Rat anti-Mouse | BioLegend | 564279 |
| 561 | 780/60 | CD14 | PE-Cy7 | Rat anti-Mouse | BioLegend | 123315 |

### Tumor specimens from BioOptions

| Inventory number | Disease category | Histo cell 1 | Histo cell 2 | Histo cell 3 | Group Stage | Gender | Age | Collection Data |
| --- | --- | --- | --- | --- | --- | --- | --- | --- |
| SB-10195 | Respiratory Lung | Carcinoma | Adenosquamous |  | IA | M | 69 | 12/1/2014 |
| NP-17864 | Male GU: Prostate | Carcinoma | Adenocarcinoma |  | IIB | M | 61 | 12/1/2014 |
| NP-16357 | Pancreas: Pancreas | Carcinoma | Adenocarcinoma | Ductal | IIB | F | 71 | 11/11/2011 |
| SB-10211 | Kidney | Carcinoma | Renal cell |  | III | F | 54 | 6/1/1998 |
| NP-10758 | GI:Stomach | Carcinoma | Adenocarcinoma | Intestinal type | IB | M | 66 | 7/26/2002 |

### Cell fractionation

We treated HCT-15 cells with 500 nM of pp242 every 6 hours for 24 hours unless otherwise stated. Control cells were grown under normal growth conditions. Experiments were performed in duplicate using 10 cm dishes. At least three 10 cm dishes were used for the cytomatrix isolation experiment. The detailed procedure of cell fractionation into the cytosol, cytomatrix and core nucleus is described previously (Shaiken et al., 2023). Briefly, cells were lysed in an appropriate volume (0.1 ml per 100 mm dish) of Buffer A: 40 mM HEPES pH 7.4, 120 mM KCl, 0.5% Glycerol, and 0.5% NP-40 with protease and phosphatase (when necessary) inhibitors and rotated for 30 min at 4°C. Lysates were centrifuged at 500 × g for 5 min, and the supernatants were collected as the cytosol. The pellets were gently washed with two volumes of Buffer A without detergent, and resuspended in Buffer B: 10 mM Tris-HCl pH 7.4, 1.5 mM KCl, 0.5% Triton X-100, 0.5% sodium deoxycholate, 2.5 mM MgCl2, 0.2 M LiCl, and protease inhibitors by rotating for 45 min and centrifuging at 2000 × g for 5 min. The supernatants were collected as the CMX. The pellets were washed with two volumes of Buffer B. The nuclear pellet was dissolved in 8M urea. Cytosol, CMX, and nuclear fractions were clarified by centrifugation at 10,000 × g for 10 min. The protein concentration was determined by Lowry. For RNA extraction and polysome profiling, all buffers were prepared in RNase-free water with the same buffer compositions and the addition of 500 U/ml RNasin. All procedures were performed at 4°C.

### Malignant cell isolation from flash-frozen tumors

Osteosarcoma and Ewing sarcoma tumors, stored at –80°C, were obtained from Texas Children’s Hospital (Jason T. Yustein lab). Human tumors were obtained from BioOptions (Brea, CA) Biorepository. Samples ranging in weight from 20 mg to 3 g were tested to isolate malignant cells from whole tissue of mouse and human tumors. Briefly, tumors stored at –80°C were cut, weighed and placed into tubes for processing, and un-used pieces were returned to –80°C for storage. Selected tumor pieces were rehydrated in Tumor Salt Solution and separated from stroma. Isolated cells were centrifuged to pellet cells. Cells were cytospined for the microscopy and staining purposes (Patent No.: US 20,210,348,993 A1).

### LC-MS/MS Analysis

Two biological replicates were used for LC-MS/MS analysis. The cytosol and cytomatrix fractions were concentrated and digested on an S-Trap™ column (Protifi, NY) per the manufacturer’s protocol. Offline high pH STAGE peptide fractionation (15 fractions combined to 5 peptide pools) for 50 µg peptides was carried out as previously described (PMID: 30093420). LC-MS/MS analysis was carried out using a nano-LC 1200 system (Thermo Fisher Scientific, San Jose, CA) coupled to Orbitrap Lumos ETD mass spectrometer (Thermo Fisher Scientific, San Jose, CA). The peptides identified from the Mascot result file were validated with 5% false discovery rate (FDR). The gene product inference and iBAQ-based quantification was carried out using the gpGrouper algorithm (Saltzman et al., 2018). The median normalized iBAQ values were used for data analysis. The differentially expressed proteins were calculated using the moderated t-test to calculate p values and log2 fold changes in the R package limma. The FDR corrected p value was calculated using the Benjamini-Hochberg procedure. Gene Set Enrichment Analysis (GSEA) (PMID: 16199517) was performed using the canonical pathway gene sets derived from the KEGG and Reactome pathway databases (Subramanian et al., 2005).

### Ribo-seq

The cytosol and CMX from control and treated cells were obtained as previously described and digested with micrococcal nuclease (MNase, Sigma-Aldrich, St Louis, MO) at 37°C for 30 min to obtain ribosome footprints. The reaction was stopped by the addition of 1.5 volumes of 4 M guanidine thiocyanate. Footprints were extracted by adding TRIzol and isolated by precipitating with GlycoBlue (ThermoFisher Scientific, Houston, TX) overnight (Reid et al., 2015). The footprints were treated with polynucleotide kinase (ENK) at 37°C for 30 min to reverse the phosphate position. We separated 17–37 nt long footprints from the gel under UV light. We froze gel slices at −80°C and then crushed them to extract RNA.

### Library Preparation

We used 10 ng of size-selected ribosome-protected RNA (17–37 nt) as the starting material for the QIAseq miRNA Library Prep Kit (cat #331502). RNA fragments were ligated to adapters at the 3’ and 5’ ends, reverse transcribed, and amplified. The resulting libraries were size selected for 185–191 bp fragments. The libraries were quantitated by qPCR using the Applied Biosystems ViiA7 Quantitative PCR instrument and a KAPA Library Quant Kit (p/n KK4824). All samples were pooled equimolarly and sequenced on a NextSeq 500 High Output v2.5 flowcell (Illumina p/n 20024906) using the Illumina NextSeq 500 sequencing instrument with a single-read configuration (75 bp). An average of 42 million reads per sample was sequenced. FastQ file generation was executed using Illumina’s cloud-based informatics platform, BaseSpace Sequencing Hub.

### Bulk Ribo-seq analysis

Two biological replicates of ribosomal footprint and mRNA libraries were generated and subjected to deep sequencing to obtain the Ribo-seq data sets. The sequence files generated by the Ribo-seq experiments were trimmed using Trimmomatic (v0.38)(Bolger et al., 2014), and sequence quality was assessed using FastQC (v0.11.8)(Andrews et al., 2010). Next, we used the bowtie aligner (v1.3.0) to remove rRNA sequences from the Ribo-seq experiment(Langmead et al., 2009). The rRNA sequences used in the rRNA depletion step were downloaded from Ensembl BioMart. The STAR aligner (v2.7.6a) was then used to align and quantify gene expression for the Ribo-seq sequence(Dobin et al., 2013), using the human genome build GRCh38 p13 v35. Differential gene expression of protein coding genes was evaluated using the R package edgeR(Robinson et al., 2010), with upper quartile and RUV (remove unwanted variation) normalization(Risso et al., 2014). Significance was achieved for a fold change exceeding 1.5× and an FDR-adjusted p value<0.05.

### Over-representation Analysis (ORA)

ORA was performed to detect enriched gene sets corresponding to pathways and biological processes based on differential gene expression. Using the Hallmark, KEGG, Reactome, and Gene Ontology Biological Process compendia (v7.3) and the Molecular Signature Database methodology (MSigDB) (Liberzon et al., 2011), we assessed enrichment with a hypergeometric test. Significance was achieved at an FDR-adjusted p value < 0.05.

### Immunoblotting

Total protein in the cell fractions were measured, equalized by concentration, and boiled at 95°C for 5 min in sample buffer. The samples were then resolved using a 4–15% Mini-PROTEAN TGX Precast Gel (Bio-Rad) at 100V for 100 min. The resolved proteins were transferred onto Immobilon PVDF membranes (Thermo Fisher Scientific) at 200mA for 60 min at 4°C. The membranes were blocked in Odyssey Blocking Buffer (Li-Cor) and probed with the respective primary antibodies at 1:1,000 dilution overnight at 4°C. The membranes were washed three times for 5 min with Tris-buffered saline with 0.05% Tween 20 (Sigma-Aldrich) and incubated with the secondary (Santa Cruz Biotech) Horseradish Peroxidase (HRP)-antibodies (1:20,000) for 1 hour at room temperature on the rocker. The blots were detected by LiCor Odyssey scanner.

### Flow Cytometry

Ewing sarcoma (A673, TC71) cells isolated from frozen tumors were used for flow cytometry analysis. Mouse cells were blocked with CD16/CD32, and human cells were preincubated with an Fc receptor-binding antibody. Cells were stained with corresponding fluorochrome-conjugated antibodies for 30 min on ice and washed with 1 ml of FCSS. Samples were analyzed using an LSRII flow cytometer (BD Biosciences).

### Statistical Analysis

Data are presented as the mean ± SD. One-way analysis of variance (ANOVA) was used to detect significant differences between the means of more than two independent groups using Prism 6 (GraphPad). We independently performed t-tests to confirm statistically significant differences between two groups. A p-value of < 0.05 was considered as significant.

## ACKNOWLEDGMENTS

We are grateful to Dr. David Y. Graham for thoughtful suggestions, valuable discussions and careful editing. We thank Dr. Jason T. Yustein for kindly providing frozen tumors, Marzhan Urazbayeva, Marina Bissekenova, Amantai Kurenbekov, Bakhytzhan Tastulekov for their support. We also thank Dr. Fabio Stossi, Abdol-Hossein Rezaeian, Antone R. Opekun and Alyssa P. Price for reading and editing the manuscript. This project was supported by Mass Spectrometry Proteomics Core, Genomic and RNA Profiling, and Bioinformatics Core, and Cytometry and Cell Sorting Core, Baylor College of Medicine, Houston TX, USA.

## AUTHORS’ CONTRIBUTIONS

**T.E. Shaiken:** Conceptualization, data curation, formal analysis, investigation, visualization, methodology, writing–original draft, writing–review and editing. **M. Shiam:** Investigation and editing, **Joel M. Sederstrom** and **Padmini Narayanan** investigation.

