## Supplementary figures and images for "Intracellular Cytomatrix, Immobilized Biocatalysis, Matrix Micromechanics and The Warburg Effect: Entanglement of Two Age-Old Mysteries of the Normal and Malignant Cell"

### Suplemental Figure 1

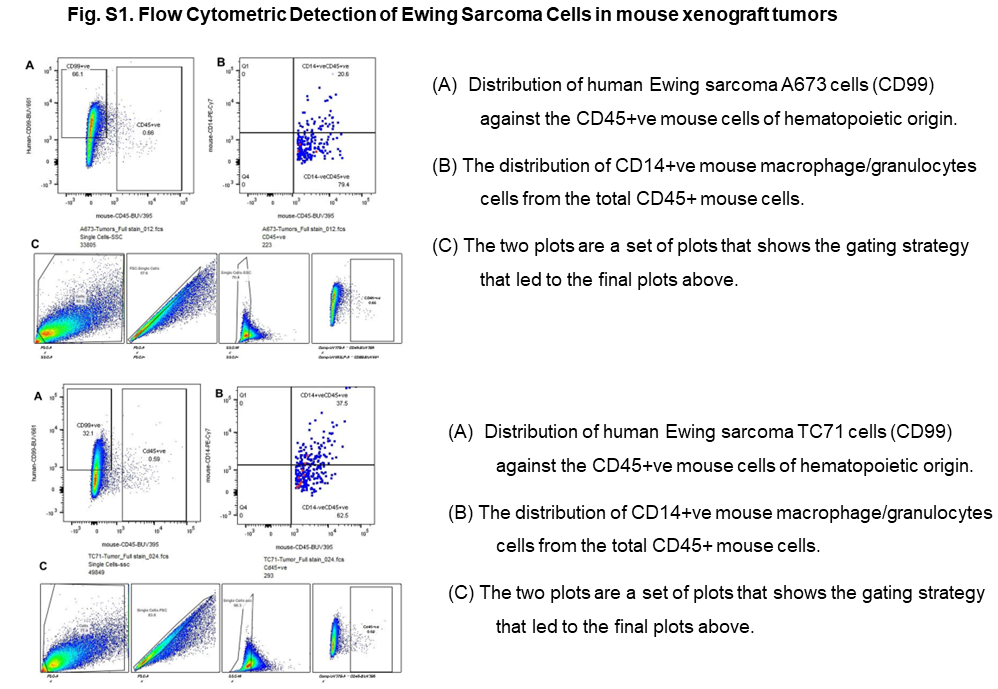
